# Cellular and transcriptomic response to pathogenic and non-pathogenic *Vibrio parahaemolyticus* strains causing acute hepatopancreatic necrosis disease (AHPND) in *Litopenaeus vannamei*

**DOI:** 10.1101/2023.11.08.566166

**Authors:** Edgar A. López-Landavery, Ángela Urquizo-Rosado, Anaid Saavedra-Flores, Sandra Tapia-Morales, Juan I. Fernandino, Eliana Zelada-Mázmela

**Author notes:** Corresponding authors: Edgar A. López-Landavery, Juan I. Fernandino, Eliana Zelada-Mázmela.

## Abstract

The shrimp industry has historically been affected by viral and bacterial diseases. One of the most recent emerging diseases is the Acute Hepatopancreatic Necrosis Disease (AHPND), which causes severe mortality. Despite its significance to both sanitation and economics, little is known about the molecular response of shrimp to this disease. Here, we present the cellular and transcriptomic responses of *Litopenaeus vannamei* exposed to two *Vibrio parahaemolyticus* strains for 98h, wherein one is non-pathogenic (VpN) and the other causes AHPND (VpP). Exposure to VpN strain resulted in minor alterations in hepatopancreas morphology, including reductions in the size of R and B cells as well as detachments of small epithelial cells from 72 h onwards. On the other hand, exposure to VpP strain is characterized by acute detachment of epithelial cell from the hepatopancreatic tubules and infiltration of hemocytes in the inter-tubular spaces. At the end of exposure, RNA-Seq analysis revealed functional enrichment in biological processes, such as the *toll3* receptor signaling pathway, apoptotic processes, and production of molecular mediators involved in the inflammatory response of shrimp exposed to VpN treatment. The biological processes identified in the VpP treatment include superoxide anion metabolism, innate immune response, antimicrobial humoral response, and *toll3* receptor signaling pathway. Furthermore, KEGG enrichment analysis revealed metabolic pathways associated with survival, cell adhesion, and reactive oxygen species, among others, for shrimp that were exposed to VpP. Our study proves the differential immune responses to two strains of *V. parahaemolyticus,* one pathogenic and the other nonpathogenic, enlarges our knowledge on the evolution of AHPND in *L. vannamei*, and uncovers unique perspectives on establishing genomic resources that may function as a groundwork for detecting probable molecular markers linked to the immune system in shrimp.

## INTRODUCTION

During the past decade, aquaculture has grown considerably as a strategic source of food supply, encompassing a diverse range of species - including fish, mollusks, crustaceans, microalgae, and macroalgae - [1]. In addition, aquaculture is the animal production sector with the fastest growth rate worldwide, even surpassing the sectors of beef, pork, and poultry [2]. The large demand for aquaculture products has resulted in heightened production levels, causing the implementation of intensive production systems and the emergence of new obstacles.

Shrimp farming is the second-largest form of aquaculture in the world, following fish production [2], and in Peru, it is the main aquaculture activity. Viral and bacterial pandemics cause high morbidity and economic losses in shrimp farming [3]. The global incidence and impact of bacterial disease on the shrimp industry have been increasing since 2010 [4], primarily due to conditions in intensive shrimp culture systems [5]. One of the bacterial diseases responsible for the collapse of shrimp production is Early Mortality Syndrome (EMS). This name was initially given because it resulted in the mortality of shrimps during the early stages of cultivation [6]; however, it was later termed Acute Hepatopancreatic Necrosis Disease (AHPND) [7, 8], because of histopathological lesions in the hepatopancreas of diseased shrimps. The first reports of AHPND with acute mortality in shrimp farming were reported in China in 2009 [8, 9], reported then in the rest of Asia [8, 10] and America [11-13], and in 2016, AHPND was added to the World Organization for Animal Health’s list of notifiable diseases [14].

AHPND results in hepatopancreatic dysfunction [15, 16], an organ that plays a crucial role in controlling systemic metabolism, digestion, and nutrient storage. Histologically, it begins with the detachment of epithelial cells from the hepatopancreatic tubules, followed by a reduction in reserve vacuoles, leading to atrophy of the hepatopancreas. Subsequently, the lesions progress from the proximal regions of the tubules to the distal areas [17]. At the most advanced stage, basophilic bacteria, which are usually opportunistic, are observed. Cell detachment persists and leads to the onset of inflammatory processes with the presence of hemocytes [18, 19]. In the terminal phase, bacterial toxins and cellular detachments cause the generalized destruction of the hepatopancreas [7]. Another vital tissue in shrimp is the hemolymph. The hemolymph carries the oxygen that is absorbed through the gills, the nutrients that are available from the digestive tract, and the catabolites that need to be disposed of through the excretory systems, hormones, and hemocytes [20]. The defensive system of crustaceans is based on cellular and humoral effectors, with hemocytes playing an essential role due to their involvement in processes such as phagocytosis, encapsulation, cytotoxicity, and nodule formation [13].

The etiological agent responsible for AHPND, known as VpAHPND, comprises a group of Gram-negative bacteria belonging to the genus *Vibrio* that naturally exist in marine and river ecosystems [21]. Their distribution largely relies on temperature, salinity, and nutrients [22, 23]. Amongst the *Vibrio* genus, *V. parahaemolyticus* is the principal vector of AHPND infection in shrimps. The strain contains a single plasmid (pVA1) of 69 kb that confers pathogenicity [24, 25] as it encodes the lethal toxins PirA and PirB, which act in a binary manner and are responsible for AHPND [26-28]. These toxins result in the detachment of cells from the epithelial tubule of the hepatopancreas (F, R and B cells) during the early stages of infection, followed by necrosis, atrophy, and massive infiltration of hemocytes. Although the histopathology of shrimp exposed to AHPND is known, the molecular response to this disease is not well understood.

Molecular techniques using PCR have been developed to identify bacteria responsible for AHPND in shrimp. These methods detect the genes for the toxins pirA, pirB, tlh, tdh, and trh in VpAHPND [27, 29, 30]. PCR methods have evolved from AP1 to AP4 [31-35]. Furthermore, omics approaches have been employed to analyze shrimp’s immune response against specific pathogens, mostly through transcriptomics. For example, researchers have employed RNA-Seq analysis to track the reaction of shrimp to *V. parahaemolyticus* challenges in organs like the hepatopancreas, hemolymph, or stomach [36, 37]. Currently, despite the development of techniques to identify the underlying cause of AHPND, our understanding of *L. vannamei*’s immune response to VpAHPND infection remains limited. In this study, we provide clear evidence of the differential response, at the cellular and molecular level, of *L. vannamei* to the challenge of two *V. parahaemolyticus* strains - one non-pathogenic (VpN) and the other pathogenic (VpP).

## MATERIALS AND METHODS

### Animal resource

Five hundred and fifty *L. vannamei* juvenile shrimps (3.0 ± 0.2 g) were collected from an intensive pond operated by Marinasol Company in Tumbes, Peru. The shrimps were transported to the Aquaculture Laboratory of the Instituto del Mar del Perú (IMARPE) with constant aeration at a density of 1 shrimp/L. The shrimps were then kept in two seawater tanks, with a water temperature of 30 ± 1°C and a salinity of 35 ups, for two weeks with constant aeration (Dissolved oxygen > 6 mg/L). Each tank contained five metric tons of seawater. They were fed with Nicovita Classic (containing 35% protein) twice daily, amounting to 4% biomass/day. Prior to being exposed to *Vibrio parahaemolyticus*, a random selection of thirty specimens were tested for the presence of the white spot syndrome virus (WSSV) disease using specific primers as recommended by the OIE [38, 39].

### Molecular characterization of *Vibrio parahaemolyticus*

#### V. parahaemolyticus origin

Three strains presumptively identified as *V. parahaemolyticus* (Vp-28, Vp-29, and Vp-32) were isolated from a semi-intensive culture of *L. vannamei* with mortality issues. These strains were obtained from the Biodes Laboratory (Tumbes, Peru). The Vp-28 and Vp-29 strains were listed as AHPND (+), which is the strain responsible for producing the symptomatology of the disease. It has the genes that express the PirA and PirB toxins (**Fig. 1a**), making it pathogenic [13]. The Vp-32 strain was classified as non-pathogenic (**Fig. 1a**). The handling of *V. parahaemolyticus* strains adhered to global biosecurity guidelines for aquatic organisms as published by the FAO.

**Fig. 1.**
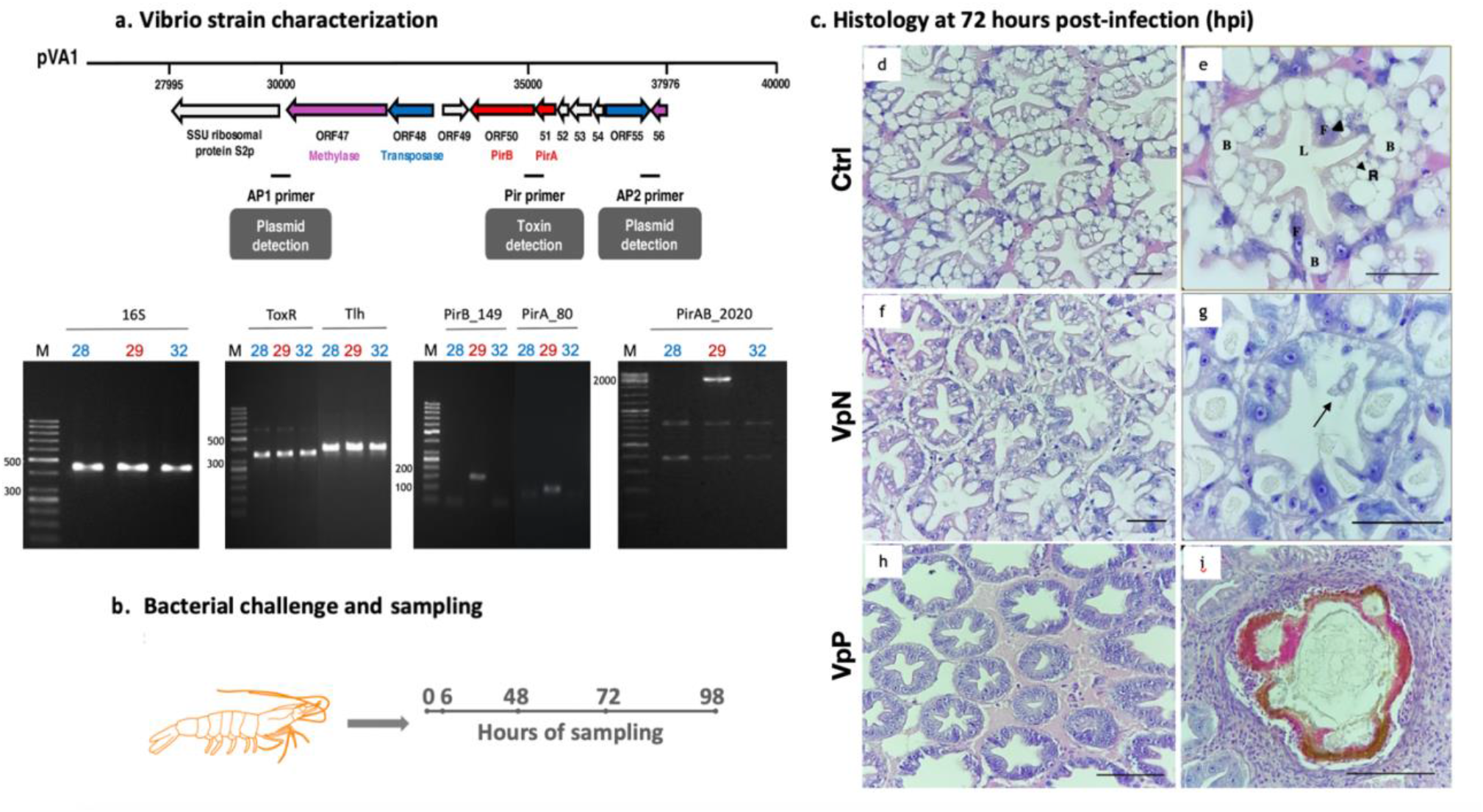
Experimental design of *Vibrio parahaemolyticus* challenge against *Litopenaeus vannamei*. a) Characterization of *Vibrio parahaemolyticus* strains. Top: Genetic structure of plasmid section containing PirA and PirB region sequences (Modified from Kumar et al., 2020). Bottom: End-point PCR for bacterial (16S), *V. parahaemolyticus* species (ToxR and Tlh), and *V. parahaemolyticus* causing AHPND (PirA and PirB) markers. b) Bacterial challenge and sampling. Shrimp *L. vannamei* was exposed to *V. parahaemolyticus* for 5 days (120 hours). Sampling for histology and RNA-Seq was conducted at 0, 6, 48, 72, and 98 hours post-infection (hpi). c) Histology at 72 hpi. Histological sections of hepatopancreas from the Ctrl (d, e), VpN (f, g), and VpP (h, i) treatments. Short arrows in e show F and R cells in hepatopancreas. Long arrow in g shows the detachment of hepatopancreatic cells to lumen. Presence of necrosis in hepatopancreatic section at image i. 28 and 32 (blue): Non-pathogenic *V. parahaemolyticus* strains, 29 (red): *V. parahaemolyticus* strain causing AHPND. Ctrl: Shrimps without *Vibrio* exposure, VpN: Shrimps exposed to *V. parahaemolyticus* lacking pirA and pirB regions, VpP: Shrimps exposed to *V. parahaemolyticus* causing AHPND. Scale bars: 100 μm.

### Bacterial culture, extraction of DNA, amplification of PCR, and Sanger sequencing

The *V. parahaemolyticus* strains obtained from the Biodes Laboratory (Tumbes, Peru) were inoculated into 40 mL of tryptic soy broth (TSB) that contained 2% NaCl and incubated overnight at 32°C. Bacterial DNA was extracted using the heat extraction method described by Phiwsaiya et al. [40]. Bacterial characterization was conducted using PCR assays that targeted the 16S rRNA gene, as well as species-specific markers for *V. parahaemolyticus* such as ToxR and Tlh, and virulence expression genes related to the pirA and pirB toxins responsible for AHPND [41-43]. The PCR reactions were carried out as described in published methods (**Table 1**). The strains’ identities were verified through Sanger sequencing, and the sequences were edited in MEGA v.7 and blasted at NCBI.

**Table 1.**
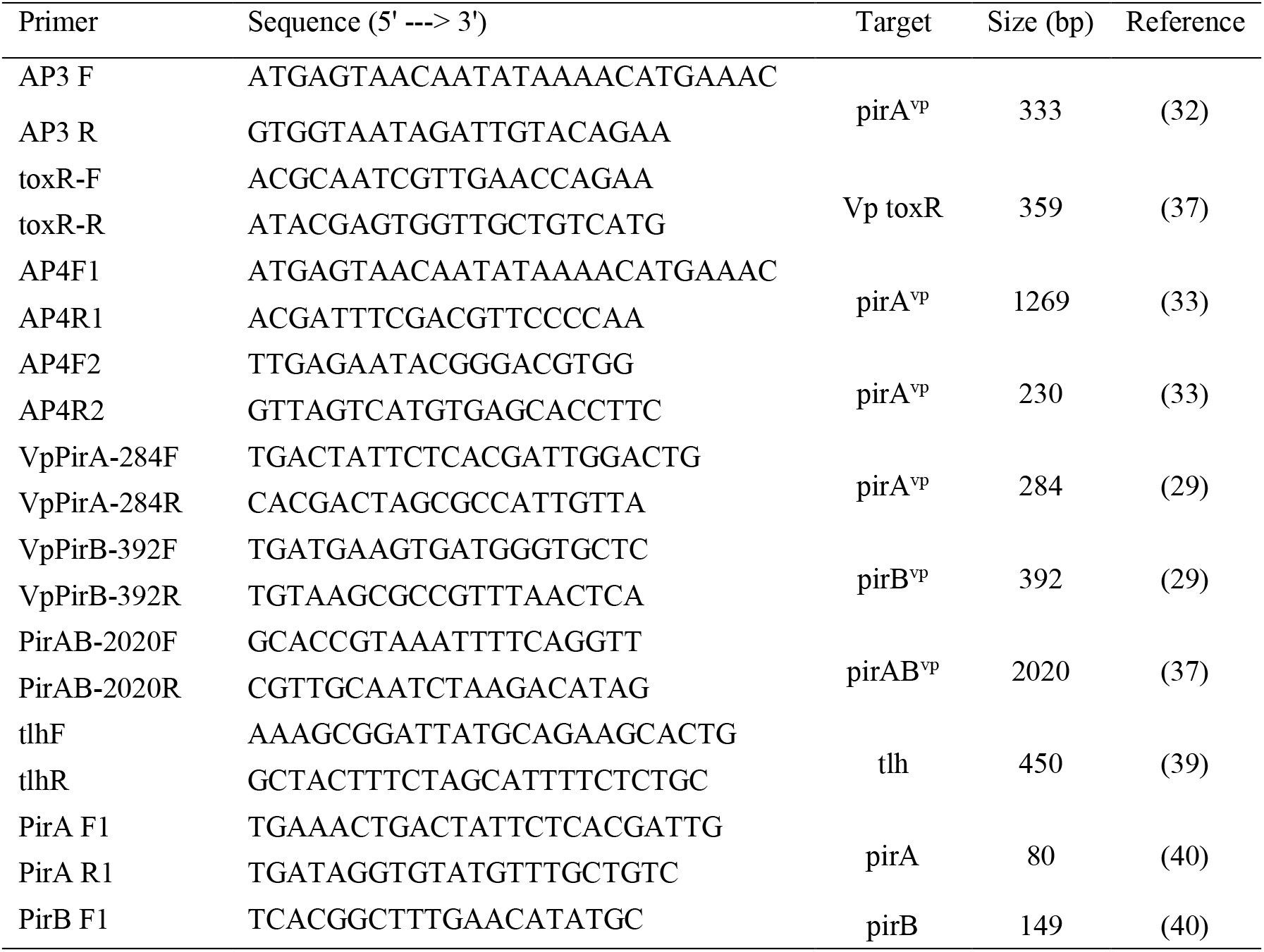
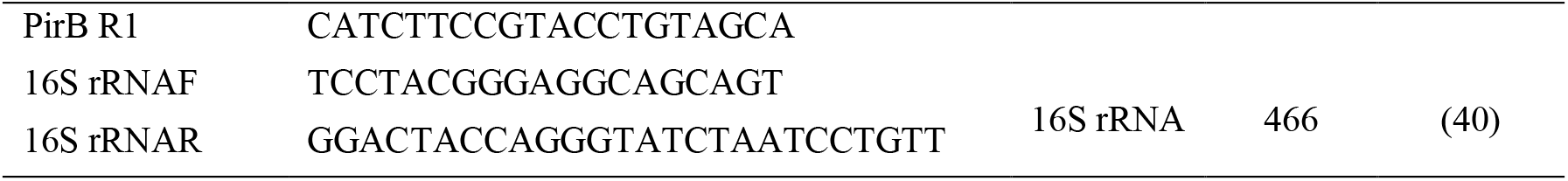
Primers sequences used to characterize *Vibrio parahaemolytivus* (Vp) strains causing AHPND. F-F2: Forward, R-R2: Reverse, toxR: toxin R, tlh: Thermolabile hemolysin.

### Bacterial challenge

#### V. *parahaemolyticus* inocula

The bacterial samples classified as AHPND (+) and non-pathogenic were suspended in 10 mL of TSB with 2% NaCl and were incubated at 32°C with continuous rotation overnight. The growth of the cells was monitored by measuring their density at 600 nm employing the NanoDrop One spectrophotometer (ThermoFisher Scientific, USA) until it reached 10^8^ - 10^9^ cells/mL. Successive dilutions were made and plated (100 μL) on Thiosulfate Citrate Bile Sucrose (TCBS, Liofilchem, Italy) agar to determine the number of colony-forming units per milliliter (CFU/mL) (data not shown). In addition, the dilutions were also plated (100 μL) on Chromatic^TM^ Vibrio (Liofilchem, Italy) for microbiological confirmation.

#### *L. vannamei* exposure to VpN and VpP treatments

The experimental design for the bacterial challenge was a completely randomized design that featured three treatments and six biological replicates, creating eighteen experimental units each with 25 shrimps. The designated treatments consisted of Ctrl (negative control without the presence of *Vibrio*), VpN (infection with non-pathogenic *V. parahaemolyticus*), and VpP (infection with *V. parahaemolyticus* that causes AHPND). Three experimental units per treatment (nine in total) were utilized solely for monitoring shrimp survival, while the remaining three experimental units were utilized for conducting histology and RNA-Seq analysis.

The bacterial challenge was adapted from Tran et al. [8], with minor adjustments. Shrimps weighing 3.5 ± 0.3 g were immersed in 30L containers filled with filtered seawater at a rate of 35 ups, exposed to UV light, and continuously aerated. For the VpN treatment, non-pathogenic strain-inoculated TSB at 2% NaCl was added; for the VpP treatment, the strain that causes AHPND was inoculated; and, for the Ctrl treatment, sterile seawater was used. For both the VpN and VpP treatments, the bacterial concentration in the experimental units was 1 x 10^6^ cells/mL. The water temperature remained consistent at 29 ± 1°C, and the shrimps were fed with Nicovita (35% protein) twice daily (3.6% biomass/day).

### Observation and sampling

The shrimps were monitored as from 6 hours post-infection (hpi) for a maximum of 5 days or until they died. For histological and transcriptomic analysis, samples of hepatopancreas and hemolymph were respectively taken from 9 shrimps per treatment at 6, 48, 72, and 98 hpi (**Fig. 1b**). Samples of hepatopancreas and hemolymph were also taken at 0h for baseline measurements. To disinfect the shrimps before sampling, a cotton swab soaked in 70% ethanol was used. The hemolymph extraction protocol was based on Wang et al. [44].

### Histopathology

The samples of shrimp hepatopancreas were fixed in Davidson’s solution for 24 hours for the histological processing. They were kept in 70% ethanol until being processed. The histological process involved dehydration, rinsing, and embedding the samples in paraffin [45]. The paraffin blocks were sliced into 5 to 6 μm sections using a Microm HM 325 microtome (Thermo Scientific), and the sections were stained with Hematoxylin and Eosin (H&E). The sections were examined using light microscopy at magnifications of 10×, 40×, and 100× to detect abnormalities, such as lesions caused by AHPND exposure. These are characterized by the massive detachment of epithelial cells from the tubules of the hepatopancreas [8]. The phases of AHPND were classified following the clinical and pathological criteria set by Soto et al. [19]. Finally, the severity of the lesions was classified according to the G classification system [46] adapted for AHPND as described by Aguilar-Rendón et al. [47]. Tissues classified as G0 had no lesions associated with AHPND, while G1 had mild focal lesions characteristic of AHPND. G2 had light to moderate multifocal lesions of AHPND, G3 had moderate to severe lesions locally extensive multifocal, and G4 had severe, multifocal to diffuse lesions.

### Transcriptomic analysis

#### Total RNA extraction and DNase treatment

The hemolymph samples of the shrimps under various treatments were taken at baseline and at 6, 48, 72, and 98 hpi for RNA extraction. Hemolymph samples were fixed in TRIzol reagent (Invitrogen) and stored at -80°C until total RNA purification. RNA was extracted following the TRIzol reagent protocol, based on the manufacturer’s guidelines. The purified total RNA was eluted with 50 μL of elution buffer and quantified by absorbance using an Epoch spectrophotometer (BioTek, USA). Total RNA integrity was assessed by 1% agarose gel electrophoresis using 1× TAE buffer prepared with DEPC water.

Three μg of total RNA from each sample were treated with DNase I (Thermo Scientific) to remove any residual DNA present in the samples. The total RNA was quantified using a Qubit fluorometer® 3.0 and the Qubit^TM^ RNA Broad Range assay (Life Technologies, USA). RNA quality was verified using the Fragment Analyzer (Advanced Analytical) and the RNA kit (DNF-471, Agilent Technologies, Wilmington, DE, USA). All samples with an RNA Quality Number (RQN) > 8 were eligible for library preparation.

#### Preparation of libraries and sequencing of RNA

To create pools of each treatment (Ctrl, VpN, and VpP) based on the infection times (0, 6, 48, 72, and 98 hpi), three hemolymph samples were collected from different individuals for each pool, with each pool representing a biological replicate. Three biological replicates were used for sequencing. Library preparation started with 1 μg of total purified RNA from each pool (Ctrl, VpN, and VpP). Libraries were prepared using the TruSeq® RNA Sample Prep v2 kit with the IDT for Illumina DNA/RNA UD indexes (Illumina, San Diego, CA, USA). Purification and fragmentation of mRNA, synthesis of cDNA, end repair, adenylation, adapter ligation, PCR amplification, library validation, equimolar normalization, and clustering were performed using the TruSeq RNA-Seq Sample Prep v2 Guide (Illumina, Part # 15026495 Rev F) using the Low Sample Protocol. Sequencing was performed on the Illumina NextSeq500 platform using the NextSeq500 System High-Output kit to generate paired reads of 74 bp (pair-end 2 x 74).

### Bioinformatic analysis

#### *De novo* transcriptome assembly

Prior to preprocessing the raw reads, their quality in fastq format was assessed using FastQC v0.11.8 [48] and subsequently consolidated using MultiQC v1.7 [49]. First, the ribosomal sequences were removed using SortMeRNA v4.3.4 [50]. Next, the reads were preprocessed using Trimmomatic v0.39 [51]. In this stage, low quality adapters and sequences with ambiguous ‘N’ bases and quality values < 30 and less than 36 bp were removed. The TruSeq3-PE-2 file, which contains the different adapters used by Illumina was used to remove the remaining sequencing adapters. Preprocessed reads were assembled *de novo* using Trinity v2.12.0 [52]. The quality of the assembled transcriptome was assessed using Transrate v1.0.3 [53] and Trinity utilities, and their completeness was assessed using BUSCO v4.1.2 [54] based on the metazoa_odb10 database.

### Transcript abundance and differential expression analysis

The resulting transcripts from the assembled *de novo* and corrected transcriptomes and Trimmomatic cleaned preprocessed reads were used to calculate transcript abundance from biological replicates. All reads were aligned against the indexed transcriptome using Bowtie2 v2.3.5 [55]. Abundance was calculated using the RNA-Seq by Expectation-Maximization algorithm (RSEM) [56] included in the Trinity pipeline. The abundance matrices from biological replicates and edgeR v3.28.1 [57] were used for differential expression analysis. The dispersion value was calculated, and then a multiplicity correction was applied to control the expected false discovery rate (FDR) using the Benjamini-Hochberg method (58). Transcripts with an adjusted *P-value* (FDR) < 0.05 and at least a two-fold change higher than or equal to 2 (FC >= |2|) were considered differentially expressed in the pair-wise comparison of the samples. Finally, Venn diagrams generated by the VENNY program were used to compare the list of differentially expressed transcripts between treatments [59].

### Functional annotation and enrichment

The transcripts were functionally annotated using the NCBI-blast-2.4.0 [60] and the Swissprot/Uniprot databases [61] released in 2021_04. Hits with an e-value < E-05 were kept. Further annotations were done using Trinotate v3.0.1 [52], assigning the best BLAST result for each protein against the SwissProt/UniProt database and making predictions for PFAM domains. For the comparison of gene ontology between differentially expressed gene lists, TopGO v2.36.0 [62] identified more significant biological processes through Fisher’s exact test and correction for FDR < 0.05. Gene Ontology (GO) terms from all annotated transcripts were used as background. The enricher function from the clusterProfiler R package [63] identified enriched KEGG pathways within DEGs lists.

### Statistical analysis

To assess the survival rate, we used the Shapiro-Wilk and Levene tests to evaluate ANOVA assumptions. A two-way ANOVA with fixed factors of treatment (Ctrl, VpN, VpP) and the time (0, 6, 24, 48, 72, 98, 120 hpi) was then conducted to test the effect of treatment and time, along with the interaction term. This was followed by a Tukey’s honestly significant differences (HSD) post-hoc test. Significance was considered at *P* < 0.05. The analysis was carried out in R version 3.6.3 [64].

## RESULTS

### Characterization of *Vibrio* strains from mortality events related to AHPND

Molecular tests were conducted on three strains of *V. parahaemolyticus* (Vp-28, Vp-29, and Vp-32) to determine if they were AHPND-causing strains based on the presence of pirA/pirB genes that are characteristic of the disease (**Fig. 1a**). All three strains (100%) tested positive for PCR that targeted the 16S rRNA gene. Also, the three strains were confirmed as *V. parahaemolyticus* by PCR targeting the tlh and ToxR genes, which are specific molecular markers. Only one of the three strains (Vp-29) tested positive for amplifying the pVA1 plasmid and binary toxin genes pirA and pirB (**Fig. 1a**, **Table 2**).

**Table 2.** Characterization and classification of *Vibrio* strains based on endpoint PCR results.

| Strain | Gene target |  |  |  |  |  |  |  |
| --- | --- | --- | --- | --- | --- | --- | --- | --- |
|  | 16S rRNA | ToxR | Tlh | Ap3 | AP41 | PirAB2020 | PirA248/<br>PirB392 | PirA80/<br>PirB192 |
| Vp-28 | + | + | + | - | - | - | - | - |
| Vp-29 | + | + | + | + | + | + | + | + |
| Vp-32 | + | + | + | - | - | - | - | - |

Sequencing analysis of the Tlh gene PCR products of Vp-28, Vp-29, and Vp-32 strains exhibited 100% identity with *V. parahaemolyticus* strain FB-11, yielding consensus sequences of 402 bp (**Table 3**). The sequencing analysis of Vp-29 straińs nested PCR-AP4 amplified products showed consensus sequences that were 100% identical to those of *V. parahaemolyticus*, *Vibrio campbellii*, and *Vibrio owensii*, isolated from AHPND cases, in the GenBank database (**Table 3**). The PCR results confirm that the Vp-29 strain exhibits the pVA1 plasmid characteristic observed in the strains responsible for AHPND in shrimp, and was employed in the VpP treatment, whereas the Vp-32 strain was utilized in the VpN treatment.

**Table 3.** Bacterial sequences with significant alignments against the sequences of the Tlh gene and pVA1 plasmid.

| Species | Strain_ID | Identity (%) | E-value | Accession number |
| --- | --- | --- | --- | --- |
| <u>*tlh*</u> |  |  |  |  |
| *V. parahaemolyticus* | FB-11 | 100.00 | 0 | CP073069.1 |
| *V. parahaemolyticus* | 20151116002-3 | 99.75 | 0 | CP034306.1 |
| *V. parahaemolyticus* | 20140829008-1 | 99.75 | 0 | CP034295.1 |
| *V. parahaemolyticus* | 20140722001-1 | 99.75 | 0 | CP034290.1 |
| *V. parahaemolyticus* | FORC_018 | 99.75 | 0 | CP013827.1 |
| <u>pVA1 plasmid</u> |  |  |  |  |
| *V. parahaemolyticus* | 20140829008-1 | 100.00 | 0 | CP034294.1 |
| *V. parahaemolyticus* | 20140722001-1 | 100.00 | 0 | CP034290.1 |
| *V. parahaemolyticus* | 20160303005-1 | 100.00 | 0 | CP034299.1 |
| *V. parahaemolyticus* | MVP1 | 100.00 | 0 | CP043422.1 |
| *V. owensii* | 1700302 | 100.00 | 0 | CP033137.1 |
|  | V180403 |  |  |  |
| *V. owensii* | pVOWZ3 | 100.00 | 0 | CP033146.1 |
| *V. owensii* | V180403 | 100.00 | 0 | CP033145.1 |
| *V. campbellii* | 20130629003S01 | 100.00 | 0 | CP020077.1 |

### Histopathology from bacterial challenge

During the experiment (**Fig. 1b**), the hepatopancreas histopathology was analyzed first. The histological analysis (**Fig. 1c**) displayed the normal appearance of the hepatopancreas in the control treatment (Ctrl), where completely healthy epithelial cells of hepatopancreatic tubule and a well-defined lumen were observed (**Fig. 1d-e**). In the VpN treatment, there were minor alterations in the morphological characteristics of the hepatopancreas relative to the Ctrl treatment. These included the initial reduction in the R and B cells size in the hepatopancreatic tubules and slight detachments of the epithelial cells in some tissue area starting from 72 hpi. Nevertheless, there was no significant detachment of epithelial cells from the hepatopancreas or hemocyte infiltrations present (**Fig. 1f-g**). In contrast, VpP exhibited distinctive AHPND lesions, including acute detachment of hepatopancreatic tubule epithelial cells, hemocyte congestion within the intertubular spaces, and bacterial plaques within the hepatopancreas in some infected shrimp samples (**Fig. 1h-i**).

### Description of AHPND stages

In the VpP treatment, all three characteristic AHPND stages were identifiable through histological examination [18, 19]. At 6 hpi, we observed initial characteristics of the initial AHPND phase. These characteristics included the elongation of the hepatopancreatic tubular epithelial cells towards the interior of the lumen (tubular lumen) (**Fig. 2a**), initiation of the detachment of the epithelial cells of hepatopancreatic tubules (**Fig. 2b**), reduction in the size of hepatopancreatic cells R and B (**Fig. 2c**), and minimal hemocyte infiltration (**Fig. 2d**). At 48 and 72 hpi, lesions typical of the acute phase of AHPND were observed. These included severe detachment of tubular epithelial cells (**Fig. 2e**), significant hemocytes accumulation of as part of the inflammatory response (**Fig. 2f-g**), greater reduction of the R and B cell vacuoles (**Fig. 2h**), and necrotic cells in the tubular lumen. Finally, at 98 hpi, characteristic lesions of the terminal phase of AHPND were identified. The hepatopancreas showed severe atrophy, and there was necrosis caused by massive cellular detachments, marked hemocyte infiltration (**Fig. 2i-j**), and bacterial proliferation leading to secondary bacterial infections (**Fig. 2k-l**).

**Fig. 2.**
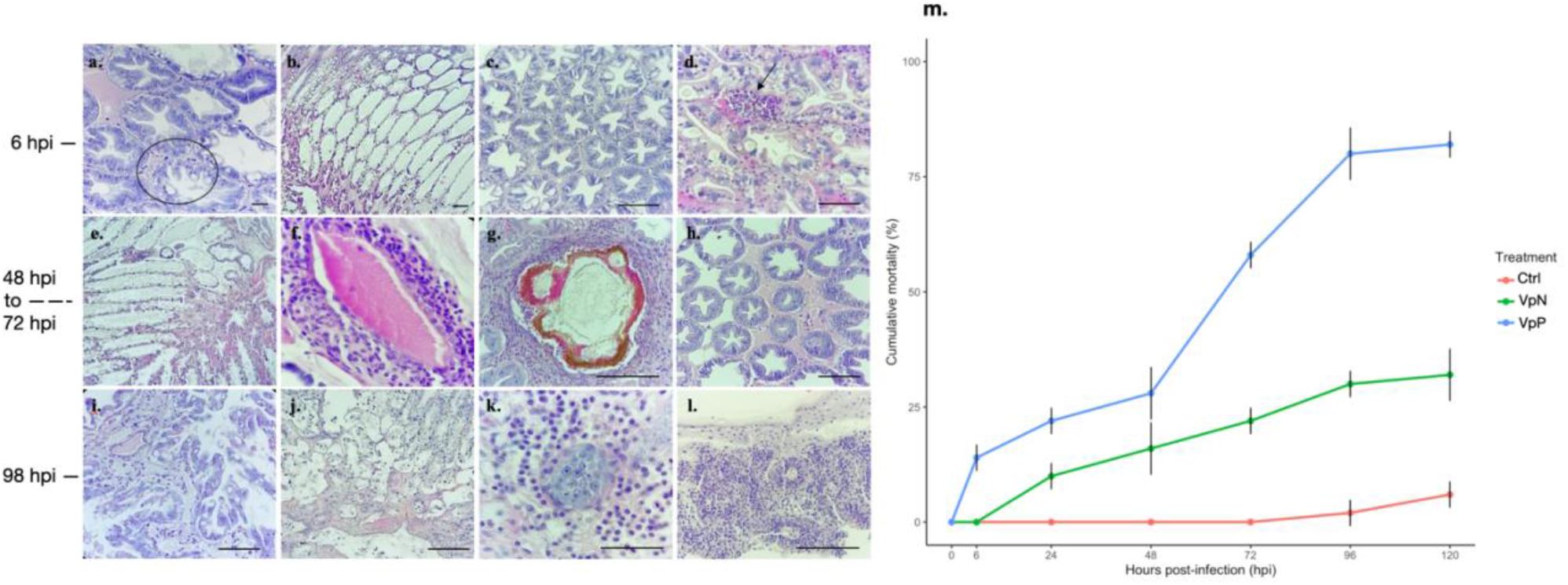
Characteristic phases of AHPND in the hepatopancreas of *Litopenaeus vannamei* corresponding to the VpP treatment. a-d) Initial stage of AHPND at 6 hpi. a) Circle shows the elongation of epithelial cells into the tubular lumen. b) Initiation of cellular detachment of the hepatopancreatic tubules. c) Decrease in the size of hepatopancreatic cells R and B. d) Arrow shows the slight hemocyte infiltration into the intertubular spaces. e-h) Acute stage of AHPND observed between 48 hpi and 72 hpi. e) Severe detachment of tubular epithelial cells and necrotic cells into the tubular lumen. f) Inflammatory response by hemocyte encapsulations of the remains of necrotic cells. g) Hemocyte encapsulations of the necrotic and melanized cell remain. h) Greater reduction of R and B cell vacuoles. i-l) Terminal stage of AHPND at 98 hpi. i-j) Disorganization and total loss of the organ’s typical structure are caused by the desquamation of the tubular epithelium. k-l) Secondary bacterial proliferation. H&E staining. Scale bars, 100 μm. m) Shrimp mortality after bacterial challenge. Ctrl: Shrimps received sterile seawater instead of bacteria, VpN: Shrimps exposed to *Vibrio parahaemolyticus* lacking pirA and pirB regions, VpP: Shrimps exposed to *V. parahaemolyticus* causing AHPND.

### Survival analysis for AHPND treatments

Monitoring of survival indicated that shrimps challenged with the Vp+ strain had a mean mortality of 82% at 120 hpi, while those challenged with the Vp- strain had a mortality rate of 32%. The Ctrl treatment exhibited a mean mortality rate of 6%. Significant mortality in the VpP, VpN, and Ctrl treatments started 6 hpi, 24 hpi, and 96 hpi, respectively (**Fig. 2m**). The two-way ANOVA revealed a significant interaction between treatment and time (*P* *** < 1.21e-12), as well as significant effects for each of the fixed factor treatments (*P* *** < 2e-16) and times (*P* *** < 5.04e-16). The Tukey’s HSD post-hoc analysis showed that the VpP and VpN treatments had significantly higher mortality rates than the Ctrl treatment (*P* *** < 0.0001). Additionally, mortality was significantly higher in the VpP treatment than in the VpN treatment (*P* *** < 0.0001).

### Transcriptomic analysis and quality assessment

The transcriptome of *L. vannamei* generated a total ∼371,200,00 pair-end raw reads (**Table S1**). Following filtration of residual rRNA with SortMeRNA v4.3.4, and removal of sequencing adapters and poor-quality reads with Trimmomatic v0.39, 342,369,271 (92.23%), clean reads were retained for assembling the *de novo* transcriptome. The initial transcriptome assembly generated 51,431 transcripts and 42,122 genes, with an average contig size of 916 pb and a contig N50 of 1,701 bp (**Table 4**). The Transrate v1.0.3 analysis reveals that 41,050 transcripts (79.82%) and 38,769 genes (92.04%) were retained, with an average contig size of 726 bp and contig N50 of 1,178 bp. In total, 29,827,713 assembled bases were identified. Bowtie2 generated an overall alignment rate of 90% between the processed reads and the assembled transcriptome. The assembly’s integrity was evaluated using BUSCO, which included 96.8% of the metazoan orthologs (**Table 4**) and showed high integrity (**Supplementary material, Fig. S1**).

**Table 4.** Statistical summary for the *de novo* transcriptome assembly and annotation from hemolymph of *Litopenaeus vannamei* under *Vibrio parahaemolyticus* challenge (Ctrl, VpN and VpP).

|  |  |
| --- | --- |
| **Final assembly** |  |
| Total Trinity genes | 38,769 |
| Total Trinity transcripts | 41,050 |
| GC (%) | 44.48 |
| *Stats based on all transcript contigs:* |  |
| Contig N50 (bp) | 1,178 |
| Median contig length (bp) | 408 |
| Average contig (bp) | 726.62 |
| Total assembled bases | 29,827,713 |
| *Stats based on only longest isoform per gene:* |  |
| Contig N50 (bp) | 1,119 |
| Median contig length (bp) | 392 |
| Average contig (bp) | 699.68 |
| Total assembled bases | 27,125,781 |
| **Assembly quality assessment** |  |
| Recovered transcripts with Transrate (%) | 79.82 |
| BUSCO completeness (%) | 96.80 |
| Bowtie2 overall alignment rate (%) | 90 |
| **Annotation** |  |
| Transcripts with coding region | 24,309 |
| Transcripts with BLAST hit (BLASTX) | 10,803 |
| Transcripts with BLAST hit (BLASTP) | 15,404 |
| Transcripts with orthologous groups (ENOG and COG) | 41 |
| Transcripts with KEGG pathway | 7,358 |
| Transcripts with GO terms | 7,081 |
| Transcripts belonging to protein families (pfam) | 8,693 |

### Functional annotation of the transcriptome

The corrected transcriptome was functionally annotated using Trinotate, as outlined in **Table 4**. Analysis of the sprot_Top_BLASTX_hits results showed a transcript annotation level of 27.87%. Additional annotations with hits or homologies were found for GO (17.25%), KEGG (17.92%), Pfam (21.18%), TmHMH (4.74%), and EggNOG (0.11%). GO clustering revealed that the most frequently occurring terms in the biological process category were related to the “regulation of transcription by RNA polymerase II”. In the molecular function category, the most prevalent terms were “metal ion binding”, “ATP binding” and “DNA binding” (**Fig. 3a**). EggNOG comprises Evolutionary Non-supervised Orthologous Groups (ENOG) and Cluster of Orthologous Groups (COG), which consists of 24 categories and transcripts. Fifteen of these were present in this study, and the most common COG categories were “posttranslational modification, protein turnover, chaperones” and “amino acid transport and metabolism” (**Fig. S2**).

**Fig. 3.**
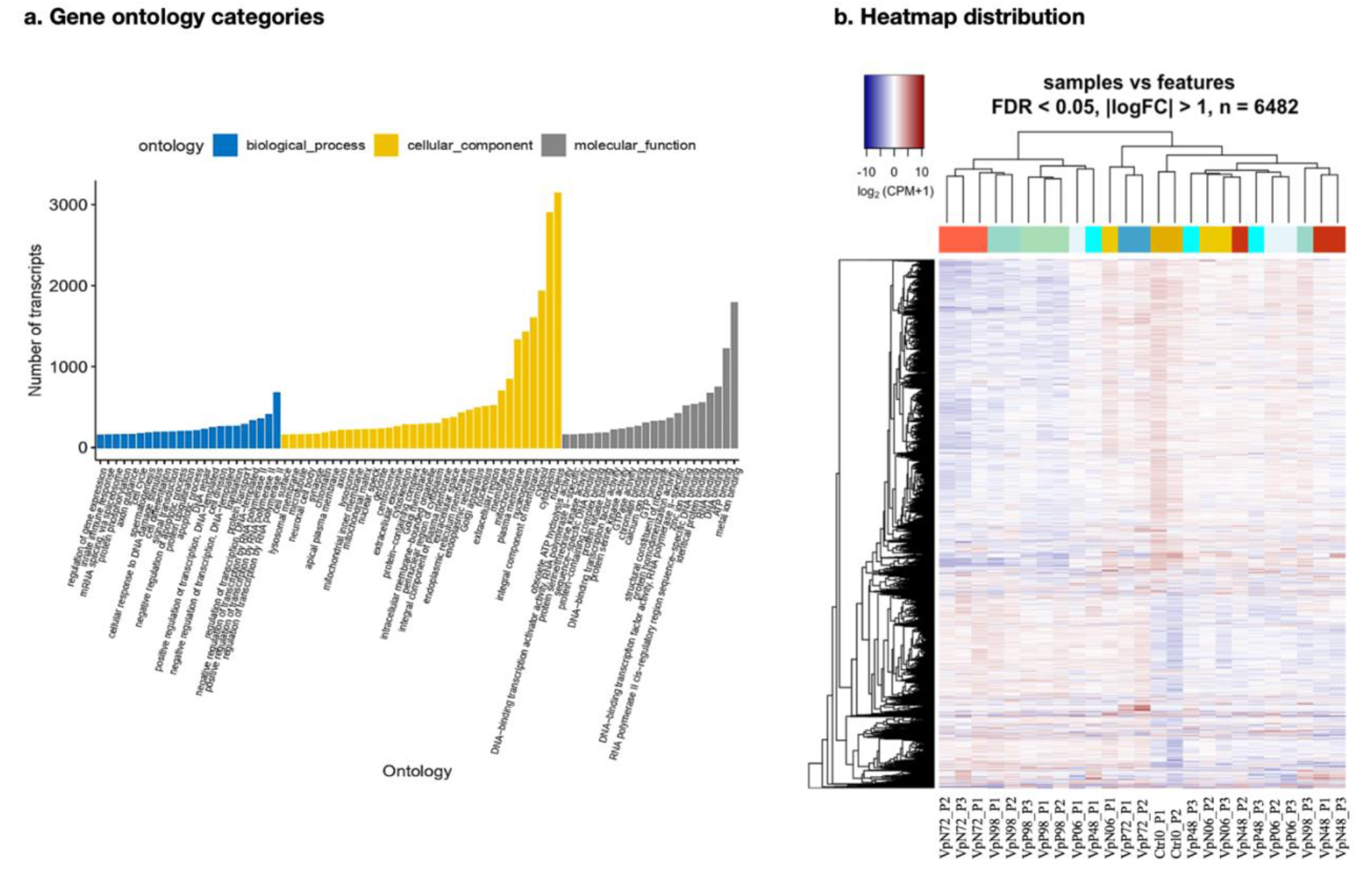
a) Distribution of Gene Ontology terms for the whole transcriptome from hemolymph of *Litopenaeus vannamei* under bacterial challenge with two strains of *Vibrio parahaemolyticus*, one causing AHPND and the other non-pathogenic, for 98 hours. b) Heatmap of the upregulated and downregulated transcripts from *L. vannamei* hemolymph under bacterial challenge with two strains of *V. parahaemolyticus*, one causing AHPND and the other non-pathogenic, for 98 hours. Ctrl: Shrimps received sterile seawater instead of bacteria, VpN: Shrimps exposed to *V. parahaemolyticus* lacking pirA and pirB regions, VpP: Shrimps exposed to *V. parahaemolyticus* causing AHPND, 0-98: Hours post-infection, P1-P3: Biological replicates. CPM: Counts per million, Red bar: Upregulated, Blue bar: Downregulated.

### Effect of bacterial challenge on differential transcriptome response

The heatmap analysis, using differentially expressed transcripts (DETs), confirmed variations in the expression patterns among treatments (**Fig. 3b**). Generally, the bacterial challenge modulated transcript up-regulation in Vp treatments when compared to the Ctrl. Biological replicates from the Ctrl treatment were clustered together. VpN and VpP biological replicates displayed similar pattern as the Ctrl, except for VpP at 48 hpi. Non-grouping at 48 hpi for VpP biological replicates may indicate differences in expression, probably as a consequence of the individual immune response.

Differential expression analysis enabled the identification of 6,482 DETs, including 5,340 corresponding to the VpN treatment (1,800 upregulated and 3,540 downregulated) and 4,342 corresponding to the VpP treatment (1,332 upregulated and 3,010 downregulated). The analysis determined that shrimps subjected to the VpN treatment had the highest amount of DETs in the hemolymph. The greatest amount of upregulated and downregulated DETs corresponded to the VpN and VpP treatments at 72 and 98 hpi, respectively (**Fig. 4a-c**).

**Fig. 4.**
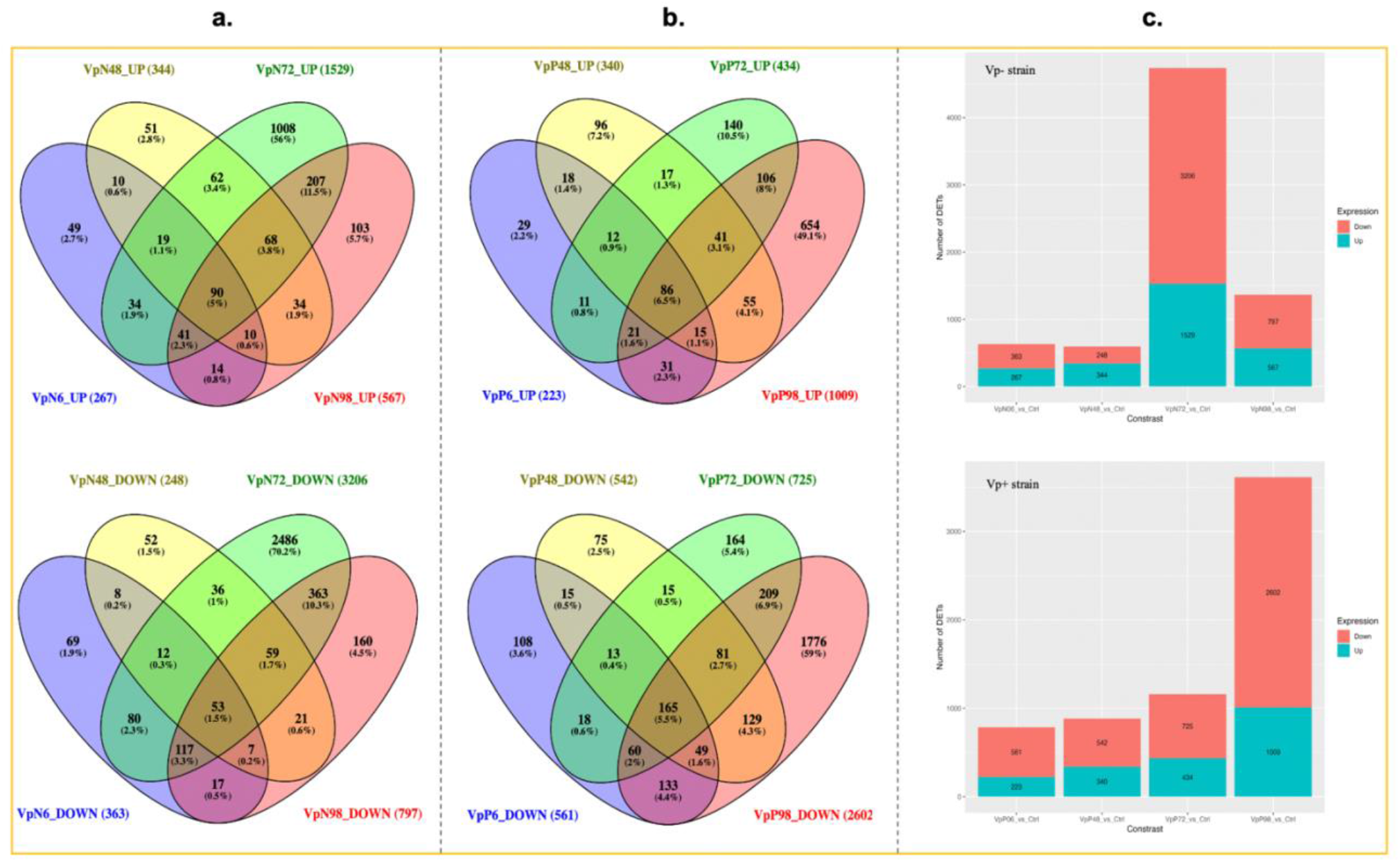
Venn diagrams for the number of differentially expressed transcripts (DETs) from hemolymph of *Litopenaeus vannamei* under bacterial challenge with two strains of *Vibrio parahaemolyticus*, one causing AHPND and the other non-pathogenic, for 98 hours. a) VpN treatment compared to control treatment, b) VpP treatment compared to control treatment, Top: Upregulated DETs, Bottom: Downregulated DETs, c) Upregulated and downregulated transcripts between contrasts using the control treatment as reference. VpN: Shrimps exposed to *V. parahaemolyticus* lacking pirA and pirB regions, VpP: Shrimps exposed to *V. parahaemolyticus* causing AHPND, 0-98: Hours post-infection, Ctrl: Control treatment.

Considering only the differential comparison between Vp treatments (VpP vs VpN), with VpN as the reference, the presence of the *V. parahaemolyticus* strain causing AHPND led to the induction of a higher number of up-regulated transcripts (734) than to down-regulated transcripts (510) in the hemolymph of *L. vannamei*. DETs were identified at 72 and 98 hpi for both up- and down-regulated cases (**Fig. 5a**).

**Fig. 5.**
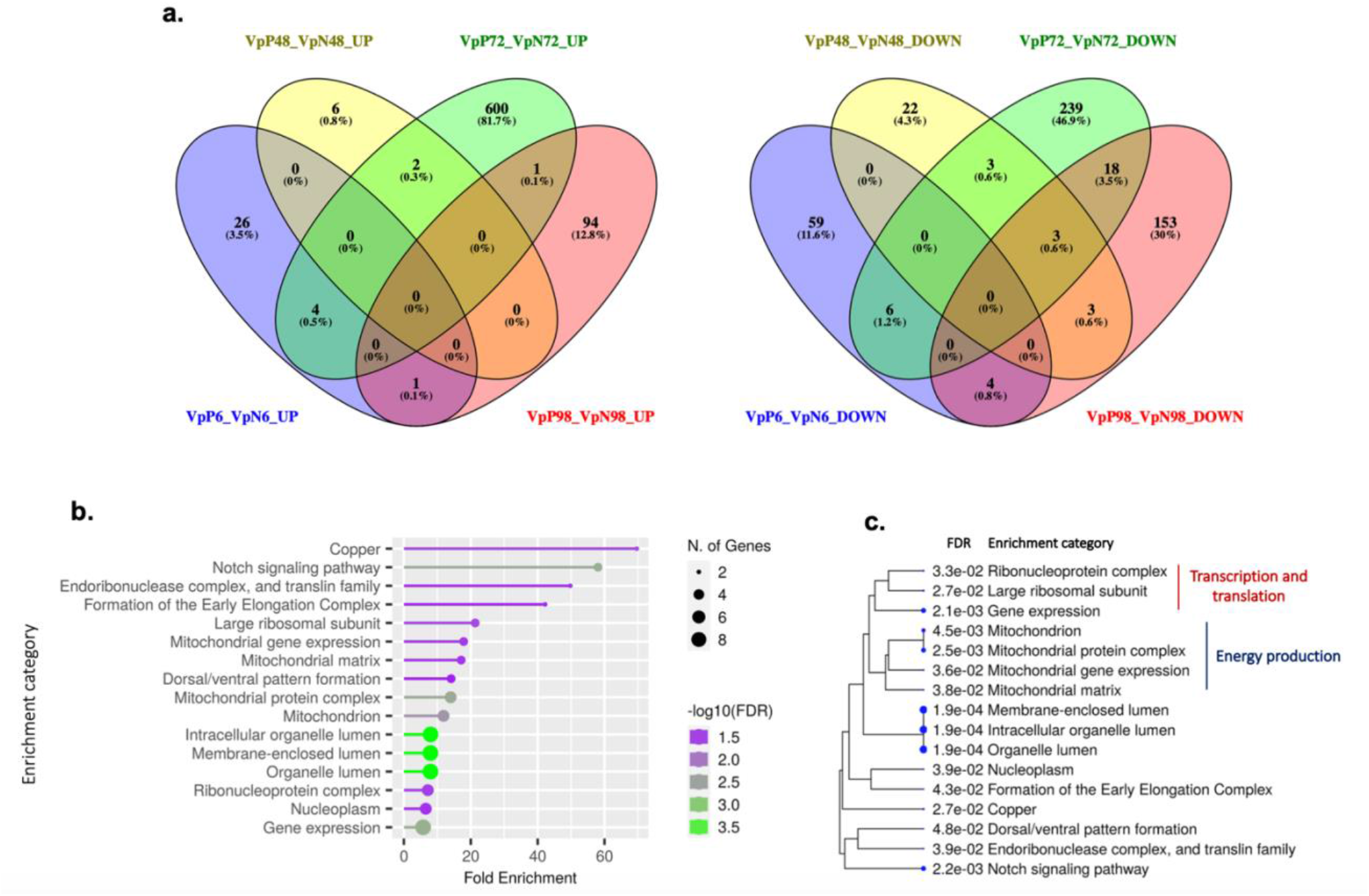
Differential expression transcripts (DET) and enrichment analysis between Vp treatments, with VpN as the reference, from the hemolymph of *Litopenaeus vannamei* under bacterial challenge. a) Up- and down-regulated number of transcripts in each comparison point. b) Enrichment categories induced by the VpP strain presence based on all DETs at 72 hpi. c) Clustering of enrichment categories and associated GO components at 72 hpi. VpN: Shrimps exposed to *V. parahaemolyticus* causing AHPND, VpP: Shrimps exposed to *V. parahaemolyticus* causing AHPND, 6-98: Hours post-infection, UP: Up-regulated DETs, DOWN: Down-regulated DETs.

### Functional enrichment analysis

#### Over-represented biological categories in the VpN treatment

A gene enrichment analysis was conducted based on DETs to identify the over-represented GO categories and KEGG pathways. The GO enrichment profile (**Table 5**) revealed biological processes at 6 hpi associated with the apoptotic signaling pathway in response to oxidative stress, the positive regulation of interferon-alpha production, deacetylation of histone H3, and upregulation of the apoptotic process in macrophages. At 48 hpi, we identified several biological processes: methylglyoxal biosynthesis, wound healing, propagation of epidermal cells, secretion of arachidonic acid, and the upregulation of hemocyte differentiation. By 72 hpi, we identified important biological processes in regulating pro-inflammatory gene expression began to be identified, including NIK/NF-kappaB signaling. In addition, this study identified several processes linked to cellular functions of the immune system, including endocytosis, virus immune response, autophagic cell death, and hemocyte proliferation. Notably, at 72 hpi, the signal transduction processes exhibited higher significance and greater abundance. Finally, at 98 hpi, biological processes related to the upregulation of cellular senescence, propagation of epidermal cells, wound healing, and inhibition of crucial proteins in various cellular signal transmission pathways were discovered (**Fig. S3**). The GO visualization was generated using REVIGO [65].

**Table 5.**
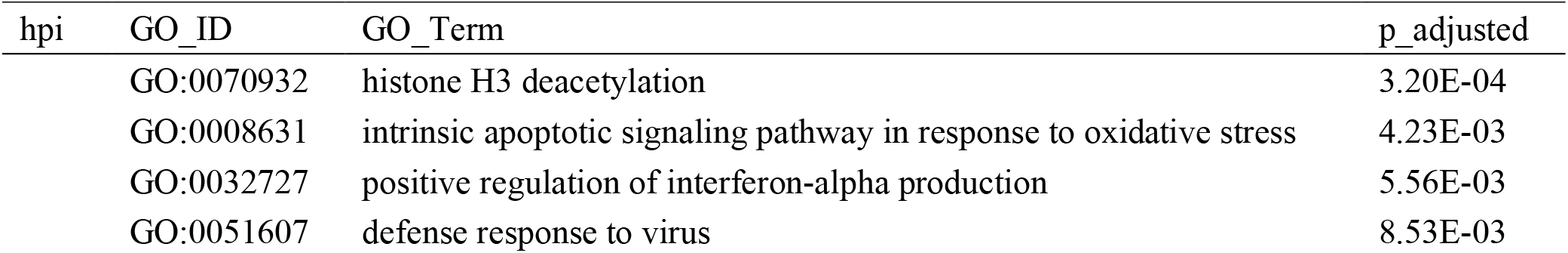
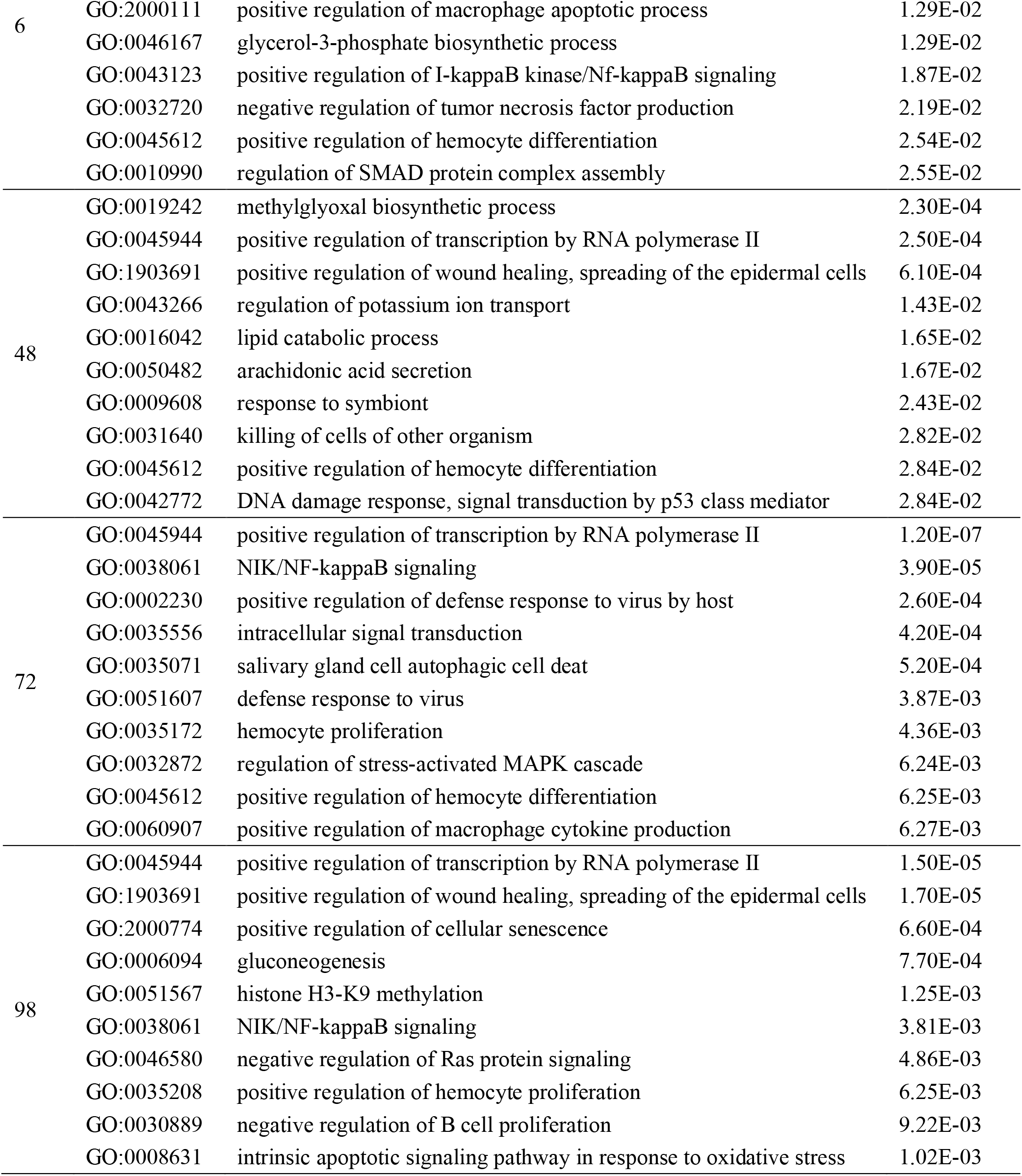
GO enrichment corresponding to up-regulated transcripts from the hemolymph of *<u>Litopenaeus vannamei</u>* <u>exposed to VpN treatment.</u>.

Regarding KEGG pathway enrichment of shrimp challenged with the Vp- bacterial strain for 98 hours, significant pathways at 6 hpi included purine metabolism, inositol phosphate metabolism, pathways associated with hormonal alterations (such as Cushing’s syndrome), and signaling pathways involved in the control of programmed cell death (Via Notch). At 48 hpi, the Ras signaling pathway was the most significantly enriched pathway, with a greater number of associated transcripts (**Fig. S5**). Other pathways related to pancreatic secretion, ether lipid metabolism, and glycerophospholipid metabolism were also identified. At 72 hpi, the Ras signaling pathway remained the most significantly enriched pathway with the highest number of associated transcripts. Furthermore, we observed an increase in the expression of pathways associated with thyroid hormone signaling, the hippo signaling pathway (a cellular communication mechanism), and MicroRNAs in cancer and shigellosis. Finally, at 98 hpi, the Ras and hippo signaling pathways were the most enriched pathways, displaying a pattern comparable to that at 72 hpi (**Fig. 6**).

**Fig. 6.**
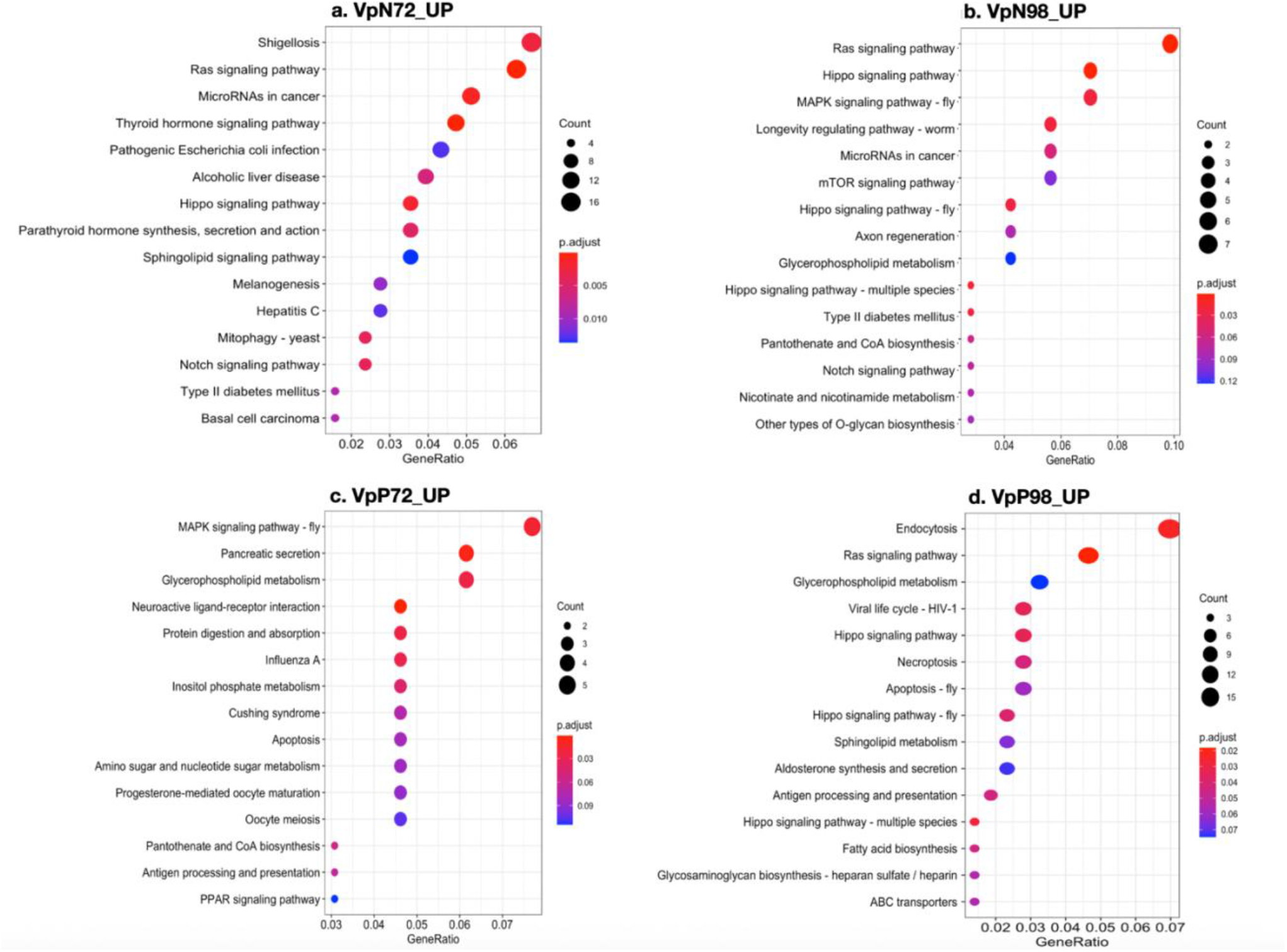
KEGG pathway enrichment based on differentially expressed transcripts (DETs) from the hemolymph of *Litopenaeus vannamei* under bacterial challenge with two strains of *Vibrio parahaemolyticus,* one non-pathogenic and other causing AHPND, for 98 hours. VpN: Shrimps exposed to *V. parahaemolyticus* lacking pirA and pirB regions, VpP: Shrimps exposed to *V. parahaemolyticus* causing AHPND. Up: Upregulated transcripts. Only 72 (left) and 98 (right) hours post-infection treatments are shown because they had a higher impact on shrimp. The control (Ctrl) treatment was used as the reference.

#### Over-represented biological categories in the VpP treatment

GO enrichment analysis (**Table 6**) of shrimp exposed to a bacterial strain causing AHPND for 98 hours revealed biological processes related to the proliferation of cells involved in the immune response, cell death triggered by the activation of T cells, biosynthetic process of coenzyme A, biosynthetic process of the glycerol-3-phosphate, and the upregulation of hemocyte differentiation. As the infection progressed (48 hpi), various molecular processes were observed, including gluconeogenesis, upregulation of transcription by RNA polymerase II, signal transduction by ARF proteins, and the biosynthesis of methylglyoxal (a substance that can trigger cell damage and activation of the immune responses), as well as apoptosis or cell death. Additionally, upregulation of hemocyte differentiation was consistently identified as a significantly enriched biological process throughout the infection process. At 72 hpi, biological processes were related to the activation of the signaling pathways of cell surface pattern recognition receptor and the upregulation of homophilic cell adhesion. Other enriched biological processes at this stage included chitin, polysaccharide, and proteolysis catabolic processes. It is important to note that enriched biological processes with the highest level of significance and abundance were identified in relation to the transport of lipids, sodium ions, and response to the bacterium. At 98 hpi, processes associated with the immune response were identified, such as positive regulation of endocytosis, defense response to the virus, positive regulation of hemocyte differentiation, viral process, and signal transduction (**Fig. S4**).

**Table 6.**
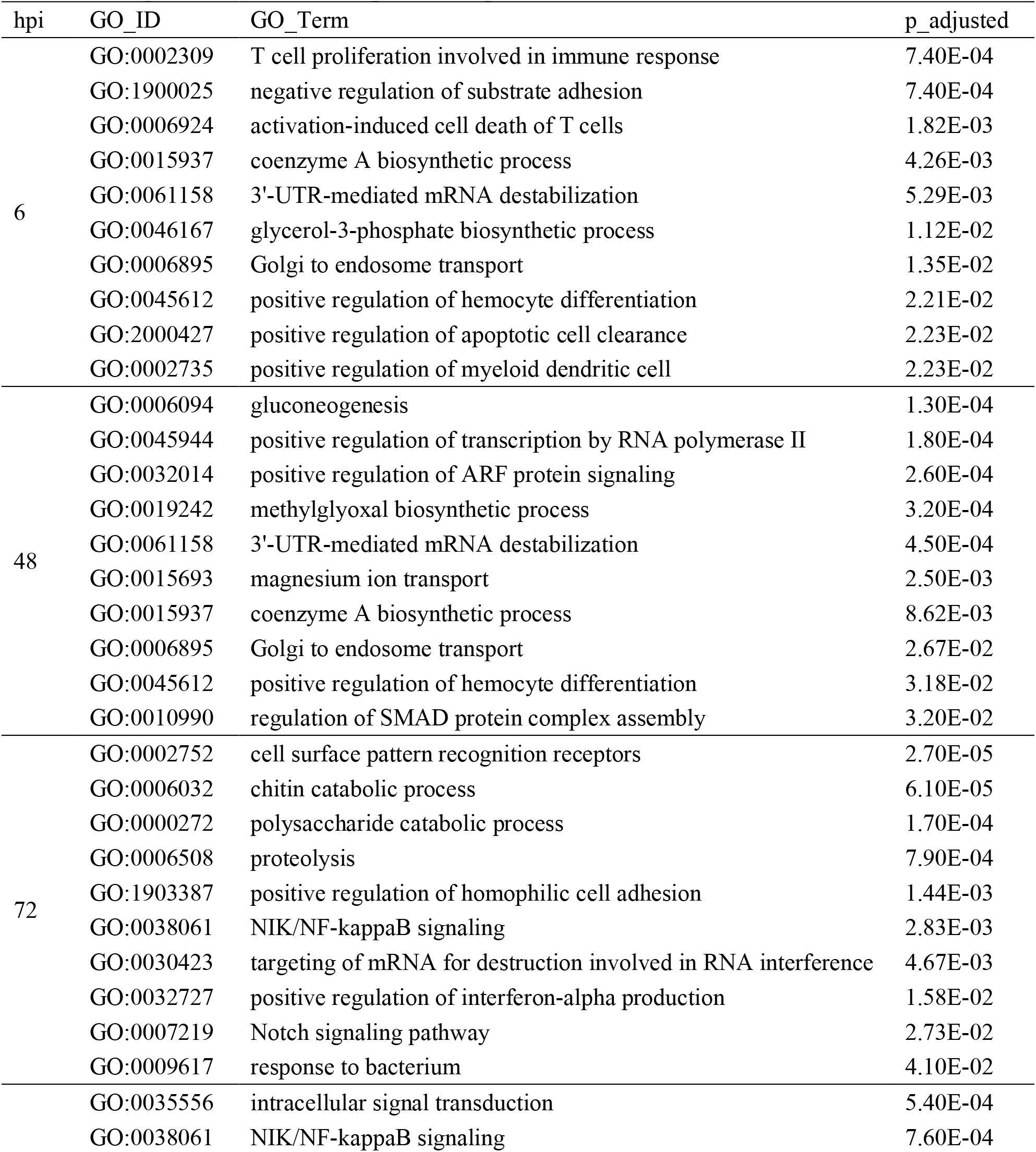
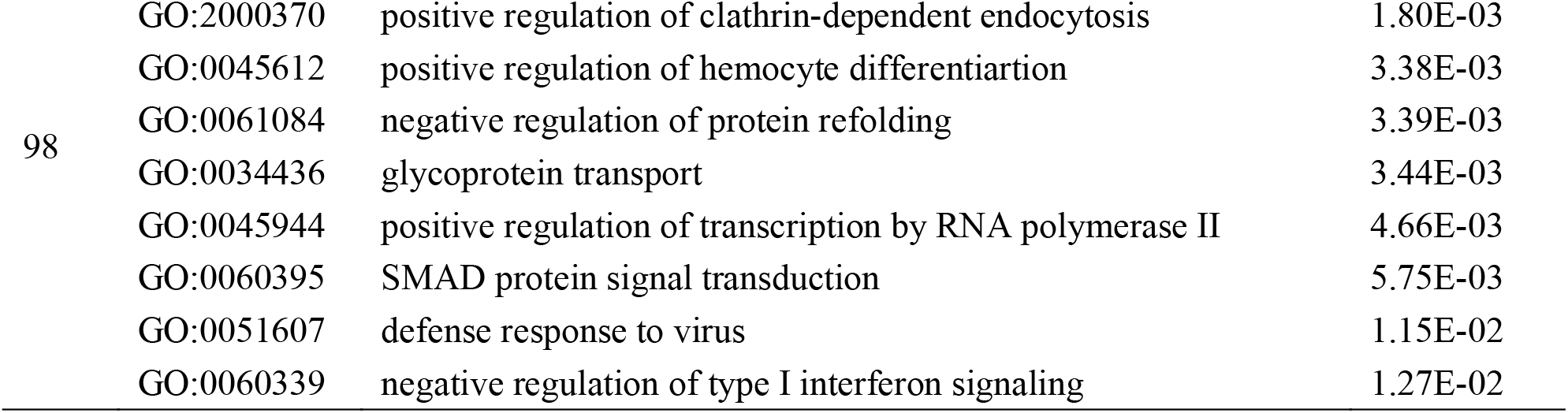
GO enrichment corresponding to up-regulated transcripts from the hemolymph of *Litopenaeus vannamei* exposed to VpP treatment.

The pathways identified through KEGG enrichment analysis at 6 hpi were involved in purine metabolism, pantothenate and CoA biosynthesis, and the synthesis and secretion of aldosterone and cortisol at 6 hpi. These pathways were also enriched at 48 hpi (**Fig. S5**). Pathways associated with the interaction of ligand-receptor neuroactive and influenza A were enriched at 48 and 72 hpi. Other enriched pathways at 72 hpi were glycerophospholipid metabolism, pancreatic secretion, and the MAPK signaling pathway. The pathways associated with Ras signaling, endocytosis, Hippo signaling pathway, viral life cycle - HIV-1, necroptosis, and antigen processing were only enriched at 98 hpi. It is worth noting that, among the mentioned pathways, the endocytosis pathway was most representative not just for its level of significance but also for the quantity of associated transcripts (**Fig. 6**).

#### Comparison between pathogenic and non-pathogenic strains (Vp treatments)

In a general context, a comparison of Vp treatments revealed the up-regulation of transcripts at 6 and 72 hpi in organisms exposed to pathogenic *Vibrio*, while the down-regulation of transcripts was observed at 98 hpi mainly (**Figs. 7, S6-S7**). The significantly enriched and overrepresented biological processes observed among Vp treatments included ribosome biogenesis (GO: 0042254), Arp2/3 complex-mediated actin nucleation (GO: 0034314), positive regulation of the Notch signaling pathway (GO: 0045747), tRNA processing (GO: 0008033), and acetyl-CoA metabolism (GO: 0006084), among others. In contrast, the underrepresented categories are related to programmed cell death (GO: 0012501), killing of cells of other organisms (GO: 0031640), defense response to bacteria (GO: 0042742), inflammatory response (GO: 0006954), necrotic cell death (GO: 0070265), muscle cell differentiation (GO: 0042692), and synaptic target recognition (GO: 0008039), among others (**Table 7**).

**Fig. 7.**
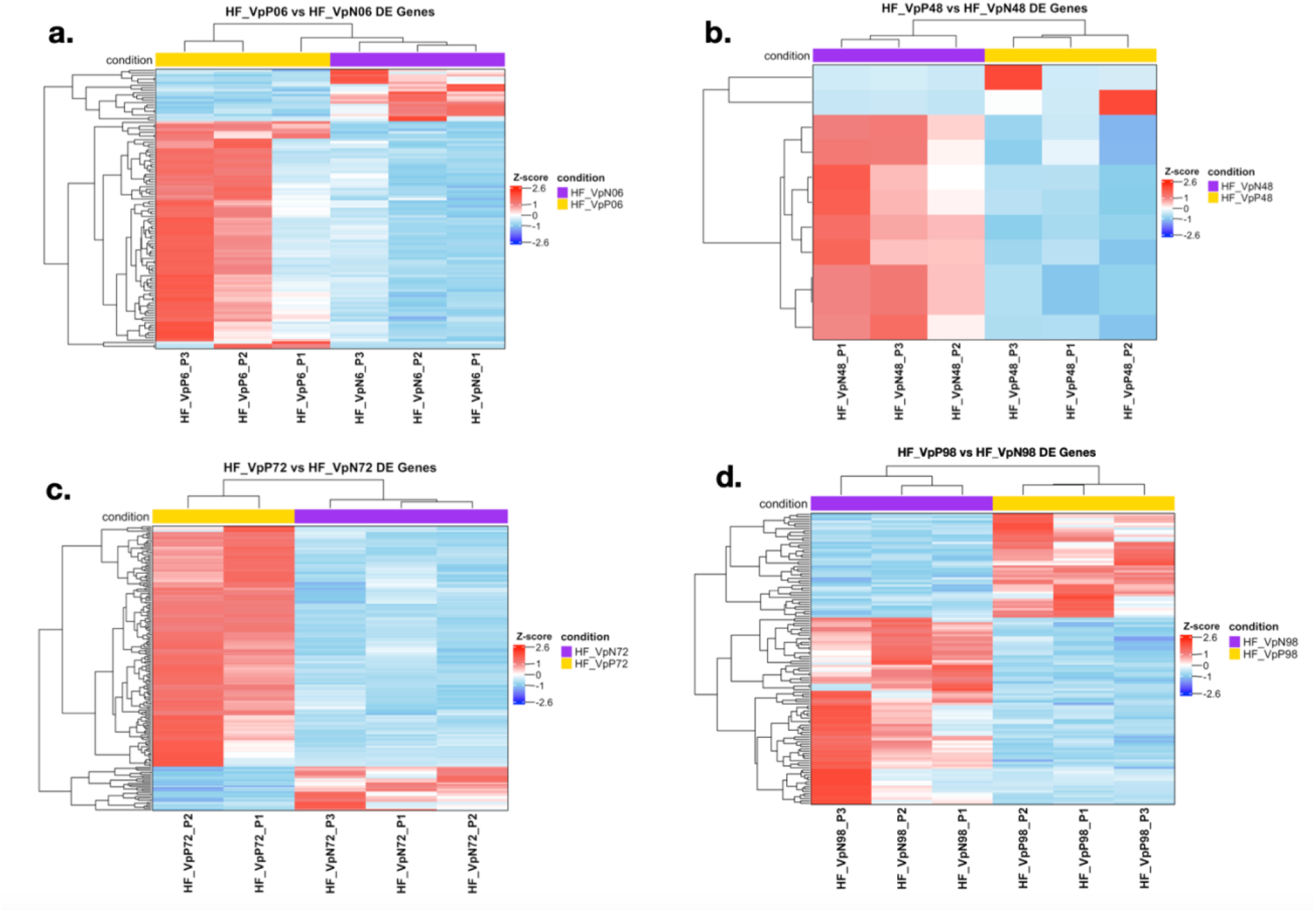
Heatmaps of the up- and down-regulated transcripts from hemolymph of *Litopenaeus vannamei* under bacterial challenge with two strains of *Vibrio parahaemolyticus*, one causing AHPND and the other non-pathogenic during 98 hours. Heatmaps represent only the comparison between the two Vp strains at each sampling point (6-98 hpi), with VpN treatment used as the reference. A) 6 hpi, B) 48 hpi, C) 72 hpi, D) 98 hpi. VpN: Shrimps exposed to *V. parahaemolyticus* lacking pirA and pirB regions, VpP: Shrimps exposed to *V. parahaemolyticus* causing AHPND, hpi: hours post-infection, red: Up-regulated transcripts, blue: Down-regulated transcripts.

**Table 7.** Top of the most significantly enriched biological processes and metabolic pathways corresponding to up- and down-regulated transcripts from the hemolymph of *Litopenaeus vannamei* exposed to VpP treatment. VpN treatment was used as the reference.

| hpi | Term | P-value | FC |
| --- | --- | --- | --- |
| <u>Up-regulated:</u> |  |  |  |
| 6 | GO:0042254~ribosome biogenesis | 3.65E-03 | 29.69 |
| 72 | GO:0034314~Arp2/3 complex-mediated actin nucleation | 9.97E-03 | 19.09 |
|  | GO:0000272~polysaccharide catabolic process | 9.97E-03 | 19.09 |
|  | GO:0045747~positive regulation of Notch signaling pathway | 1.68E-02 | 14.68 |
|  | GO:0008033~tRNA processing | 1.81E-02 | 7.07 |
|  | GO:0051791~medium-chain fatty acid metabolic process | 4.60E-02 | 42.42 |
|  | GO:0010954~positive regulation of protein processing | 4.69E-02 | 31.81 |
|  | KEGG_pathway:xla04144~Endocytosis | 3.21E-04 | 26.43 |
|  | KEGG_pathway:xla04810~Regulation of actin cytoskeleton | 4.87E-03 | 26.42 |
|  | KEGG_pathway:xla04530~Tight junction | 6.74E-03 | 22.65 |
|  | KEGG_pathway:pon03040~Spliceosome | 4.58E-02 | 35.23 |
| 98 | GO:0006084~acetyl-CoA metabolic process | 2.94E-02 | 65.98 |
|  | GO:0015677~copper ion import | 4.38E-02 | 43.99 |
|  | GO:0006085~acetyl-CoA biosynthetic process | 4.38E-02 | 43.99 |
|  | GO:0030433~ubiquitin-dependent ERAD pathway | 4.68E-02 | 7.61 |
|  | KEGG_pathway:dme00520~Amino sugar and nucleotide sugar metabolism | 4.61E-02 | 28.78 |
| <u>Down-regulated:</u> |  |  |  |
| 6 | GO:0006954~inflammatory response | 4.99E-03 | 26.39 |
|  | GO:0012501~programmed cell death | 2.36E-02 | 79.18 |
|  | GO:0045087~innate immune response | 2.62E-02 | 11.10 |
|  | GO:0046718~viral entry into host cell | 4.58E-02 | 32.99 |
| 48 | GO:0042742~defense response to bacterium | 2.72E-02 | 63.63 |
| 72 | GO:0042692~muscle cell differentiation | 7.38E-03 | 22.27 |
|  | GO:0019242~methylglyoxal biosynthetic process | 2.65E-02 | 74.23 |
|  | GO:0070265~necrotic cell death | 3.95E-02 | 49.49 |
|  | GO:0015881~creatine transport | 4.23E-02 | 37.11 |
|  | GO:0060766~negative regulation of androgen receptor signaling pathway | 4.63E-02 | 37.11 |
| 98 | GO:0050832~defense response to fungus | 5.67E-03 | 25.66 |
|  | GO:0007379~segment specification | 3.32E-02 | 58.17 |
|  | GO:0031640~killing of cells of other organism | 3.32E-02 | 58.17 |
|  | GO:0045892~negative regulation of transcription, DNA-templated | 3.52E-02 | 3.93 |
|  | GO:0008039~synaptic target recognition | 3.98E-02 | 48.48 |
|  | GO:0006357~regulation of transcription from RNA polymerase II promoter | 4.54E-02 | 2.36 |
|  | GO:0006936~muscle contraction | 4.62E-02 | 41.55 |

In a general context, a comparison of Vp treatments revealed differential over- and underrepresented biological categories. The biological processes that were overrepresented included transcription, translation, protection against reactive oxygen species (ROS), and energy generation (**Fig. 5b-c**). The biological processes that were underrepresented primarily pertained to the immune response, down-regulation of transcription, and likely inhibited growth due to a low rate of muscle cells differentiation. On the other hand, the enrichment of KEGG pathways, based on DETs, indicated a substantial number of significant pathways at 72 hpi in overrepresented categories (**Fig. 6c**). These pathways are associated with endocytosis (xla04144), regulation of actin cytoskeleton (xla04810), tight junction (xla04530), and spliceosome (pon03040) (**Table 7**).

## DISCUSSION

*Vibrios* are prevalent bacterial pathogens in shrimp farming; however, these bacteria being part of the shrimp microbiota and being typically opportunistic pathogens. Some pathogenic strains can cause diseases, such as AHPND [66], because bear a plasmid that encodes the lethal toxins pirA and pirB [21, 25, 26], which operate in a binary manner and lead to severe hepatopancreas atrophy in shrimp [28, 29]. In this study, we evaluated the cellular and transcriptomic response of *L. vannamei* through exposing the shrimps to challenge with two strains of *V. parahaemolyticus*, including one non-pathogenic (VpN) and the other pathogenic (VpP) strain.

### Histopathology and survival rate in response to Vp infection

Histopathological analysis of the hepatopancreatic tissues in *L. vannamei* provides information on morphological changes, tissue lesions, their severity, and cellular responses to the presence of VpN and VpP strains [27, 40, 67, 68]. In the present study, the control group (Ctrl) exhibited uniform structures of the hepatopancreatic tubules with healthy digestive cells E, F, R, and B, as previously reported [69-71]. These findings suggest normal nutrient assimilation and functioning of the defense system.

In VpN treatment, there were observations of digestive cell detachment and intact cells in the tubular lumen, indicating vulnerability to bacterial attack. This damage was identified in samples taken at 72 and 98 hpi, with mild (G1) to moderate (G2) levels. These specialized cells become detached from the hepatopancreatic tubules due to their fragility and susceptibility to attacks by pathogens [72]. However, these cell detachments did not correspond to lesions that are characteristic of AHPND, as previously described [6, 16, 73]. Additionally, there were no hemocytic infiltrations observed in the intertubular spaces of the hepatopancreas. This absence of infiltration may be attributed to a less aggressive bacterial attack, that did not elicit a strong response from the defense system [76, 77]. A decrease in vacuole size within R and B cells at 72 hpi indicated a reduction in food consumption, which could lead to nutrient deficiency and damage to hepatopancreatic cells [78]. This damage, which primarily affects epithelial cells in the hepatopancreas [79], disrupts its digestive, metabolic, and detoxification functions [80, 81].

Shrimps challenged with VpP strain experienced more severe hepatopancreatic damage compared to VpN strains. The severity of the injury was greater in VpP treatment (G3 to G4) compared to VpN (G1 to G2). Reduction in R- and B-cell size was more pronounced in VpP, potentially due to both bacterial concentration and the release of pirA and pirB toxins. AHPND-induced lesions were more widespread and severe in the VpP treatment, potentially caused by the release of pirA and pirB toxins linked to AHPND evolution [82]. Typical sharp AHPND phase lesions were also detected in the VpP treatment at 48 and 72 hpi. Extensive (G4) harm to the tubular epithelium was occurring with conspicuous cell detachment and necrosis. This condition aligns with other studies on *L. vannamei*, indicating that during this stage of AHPND, inflammatory processes begin with strong hemocytic infiltrations and potential cellular responses [9, 83]. Notably, hemocytes serve as the first line of defense in shrimp [84-86]. It is important to note that the timing of the acute AHPND phase identification in this study was not consistent with previous findings [19], which indicate that the acute AHPND phase is present in shrimps exposed to *V. parahaemolyticus* from 3 to 18 hpi. This difference may be due to differing virulence among *V. parahaemolyticus* isolates that cause AHPND, and the shrimp’s ultimate response depends on the infection dose [87] and exposure method. The terminal phase of AHPND at 98 hpi showed complete disorganization of hepatopancreatic tubules, extensive cell detachment, and hemocytic infiltrates. Secondary bacterial proliferation suggests a compromised system functioning due to loss of digestive function and nutrient assimilation. These features have been recognized as pathognomonic indicators of this phase of AHPND in various studies [8, 68] and are directly linked to the serious hepatopancreatic injury (G4) that led to the death of shrimp in this experiment.

Shrimp mortality in our study was comparatively lower than in other studies [19, 88]. This may be attributed to strain virulence, individual health status, and immune system activation [88]. Our findings suggest that the VpP strain causing AHPND may have been less virulent in comparison. Also, it is worth noting that previous reports suggest genetic lines of shrimp used in South America may have some resistance towards AHPND [89]. In the case of the VpN strain, there was mortality, but it was significantly lower than the mortality caused by its positive counterpart. Previous studies have reported contrasting mortality rates (0 vs. 75%) in shrimp exposed to strains of *Vibrio* that lack pirA and/or pirB regions [88, 90, 91]. Thus, genetic resistance and the absence of specific virulence factors appear to have contributed to the variable mortality rates among different Vp strains. Our study provides insight into the intricate relationship between bacterial virulence, shrimp health, and immune responses in the progression of AHPND, suggesting additional genomic analysis to uncover unreported virulence factors.

### Transcriptomic response

#### Comparison between VpP strain and the Ctrl treatment

The enrichment analysis of KEGG pathways in the VpP treatment identified a responsive immune system. At 6 hpi, the most enriched pathway was observed in pathways associated with hormonal disorders triggered by high cortisol concentrations [92, 93]. Cortisol, widely known for its role in stress response and immune modulation, was implicated in various homeostatic maintenance, anti-inflammatory, and immune system functions [94, 95]. In crustaceans, the release of hyperglycemic hormone (CHH) can be triggered by stressors such as hypoxic conditions, high temperatures, and parasitic infections [96, 97]. It has been found to have effects similar to those of cortisol. Enriched pathways at 48 and 72 hpi suggested the involvement of the neuroactive ligand-receptor interaction and the MAPK signaling pathway. The latter is activated in response to both environmental and cellular stressors as well as pathogen infections [103-105]. The immune response of this pathway has been observed in shrimp following infection by *V. parahaemolyticus* [106], *V. alginolyticus* [107], *Eriocheir sinensis* [98], and *L. vannamei* [99]. The histopathology observed suggests that the MAKP signaling pathway plays an important role in transducing immunological signals in the hemolymph of shrimps in response to microbial invasions. This pathway promotes both hemocyte proliferation and differentiation, as suggested previously [108]. Genes such as Trypsin and Prostaglandin E2 (PGE2) are components of the neuroactive ligand-receptor interaction and were upregulated, indicating their significance in immune responses [100-102]. In this context, the enrichment of neuroactive ligand-receptor interaction would be a response to failures in the digestive process in the hepatopancreas and its need for greater production of circulating hemocytes due to the damage generated by the VpP strain at the acute AHPND phase.

At the end of the bacterial challenge, at 98 hpi, there was an observed increase in endocytosis and Ras signaling pathways. These pathways, including Ras, Raf, and MAPK pathways are responsible for transmitting signals from the extracellular environment to the nucleus to activate genes related to cell growth and differentiation [109], as well as wound healing, tissue repair, and cell migration [110, 111]. In crustaceans, the Ras signaling pathway activates and regulates the phagocytosis performed by hemocytes [112]. This is consistent with the progression of hepatopancreatic lesions based on the histology from 48-72 hpi to 98 hpi, which is characterized by strong hemocyte migration, wounds, nodules, melanization, and tissue disorganization. In other penaeids, like *Fenneropenaeus chinensis*, studies have reported upregulation of various genes linked with endocytosis (which the Ras pathway activates) due to WSSV infection [113]. Therefore, our findings propose that prolonged exposure to the VpP treatment leads to the upregulation of genes associated with the Ras signaling pathway, facilitating the elimination of the VpP strain through endocytosis.

#### Comparison between VpP and VpN strains

Comparing both Vp strains, the biological processes and metabolic pathways that were enriched showed contrasting natures. Shrimp exposed to the VpP strain initially induced a high rate of ribosome biogenesis, likely associated with the high demand generated by the translational process. At 72 hpi, pathways related to *Vibrio* invasion and dispersal, along with energy generation through carbohydrate, lipid, and protein catabolism, were enriched. The process of ribosome biogenesis is a highly intricate and energy-intensive process. This suggests a need for fast activation of the shrimp translation machinery against the VpP strain. Prior studies on *L. vannamei* have reported increased ribosome biogenesis in individuals exposed to AHPND and aflatoxins [115, 116].

As the disease progressed, tight junctions (TJ) were induced, likely as a defense mechanism to prevent pathogen entry. TJ disruption has been linked with the process of releasing toxins pirA and pirB, as well as bacteria, from the stomach to the hepatopancreas during the AHPND infection process in marine shrimps [118, 119]. Based on the histology, it is possible that the induction of TJs in this study is linked to the requirement for increased production to obstruct the entry of toxins or *V. parahaemolyticus* into the hepatopancreas. Additional enriched categories at 72 hpi were associated with the utilization and synthesis of energetic substrates such as carbohydrates, lipids, and proteins, possibly signifying the need to maintain homeostasis and prevent the collapse of hepatopancreatic tissue. This indicates a collaborative endeavor to uphold tissue equilibrium during infection. Pathways associated with high energy requirements, such as the degradation of misfolded proteins through the ubiquitin-dependent ERAD pathway and copper ion utilization, surfaced at 98 hpi, which coincided with the necessity for oxygen transportation via hemocyanin. The induction of acetyl-CoA processes at 98 hpi suggests that shrimp utilized other substrates as energy sources, such as alanine, glycerol, and lactate, through gluconeogenesis [122]. The above enriched biological processes and metabolic pathways observed, particularly at 72 and 98 hpi, indicate a disturbance in the homeostatic state of the shrimps at the end of the bacterial challenge. These findings suggest that the VpP strain has a significant detrimental effect on the shrimps.

### Differentially expressed transcripts are involved in the immune response

Differential gene expression sheds light on the immune response mechanisms (**Tables S2**-**S3**). The increase in expression of low-density lipoprotein receptor (LRP1)-related protein 1 indicates its involvement in pathogen recognition and immune regulation [123]. The activation of the MAPK and Toll pathways is in accordance with previous research, highlighting their importance in immune responses [124-127]. On the other hand, antimicrobial peptides (AMPs) played a crucial role in both strains, especially anti-lipopolysaccharide factors (ALFs) and lysozymes, which contribute to bacterial [130] and fungal [131] defenses. ALFs are effector molecules of innate immunity in arthropods, capable of lipopolysaccharide binding and pathogen neutralization activities [132]. After exposure to bacterial infections, shrimps show significantly increased expression of ALFs [133]. Another type of AMP upregulated in the VpP treatment was lysozyme. This enzyme is linked to defense mechanisms and digestive system [134, 135]. The interaction between these pathways and gene expressions highlights that shrimp defense against *V. parahaemolyticus* engages multiple layers of immune response, with the Toll signaling pathway playing a crucial role in shrimp’s innate immune response to *V. parahaemolyticus*, predominantly in the final phase of infection. Furthermore, the upregulation of several genes associated with apoptosis process in the hemolymph of shrimp infected by the VpP strain may be related not only to the straińs level of pathogenicity but also to the damage level caused by it in shrimp hepatopancreatic cells, which are considered energy promoters.

It is known that apoptosis is a programmed mechanism utilized not only in response to the elimination of pathogens but also to remove damaged cells, ensuring proper development and homeostasis of neighboring cells [137]. Research on marine organisms has reported upregulation of beclin 1 in the presence of toxic substances or pathogenic microorganisms, signifying its participation in the host’s immune response [140, 141]. Furthermore, a greater number of upregulated genes were linked to apoptosis [142-144] in shrimp under the VpP strain challenge, including apoptosis-resistant E3 ubiquitin-protein ligase 1, caspase-8, caspase-14, and cell death protein 3, among others. The upregulation of caspases and cell death protein 3 only in shrimp challenged with the VpP strain with time modulation at 72 and 98 hpi, suggests higher damage at the cellular level based on histology. Therefore, it is reasonable to infer that a larger quantity of proteins related to this process could be triggered to regulate this kind of event.

## CONCLUSION

This study’s functional enrichment analysis revealed the complex immune response of *L. vannamei* to *V. parahaemolyticus* infection, highlighting key pathways involved in pathogen recognition, immune activation, and cellular defense. The differing responses between VpP and VpN strains emphasized the role of virulence in influencing immune interactions. These findings enhance to our comprehension of AHPND pathogenesis and assist in designed plans for disease management and shrimp health enhancement.

## DATA AVAILABILITY STATEMENT

Data supporting this work is available at GenBank. All sequences in fastq format were deposited in the Sequencing Read Archive (SRA) of NCBI with the accession numbers from SRR19662274 to SRR19662298. The BioProject ID of our data is PRJNA849497 and the BioSample accession is SAMN29059449.

## AUTHOR CONTRIBUTIONS

EALL contributed to conception of the study. EALL, JIF designed the study. STM, AUR performed bacterial characterization. EALL, AUR performed the experiment and collected the samples. AUR, EALL performed histopathological examination. ASF, AUR performed RNA extraction and construction of RNA-Seq libraries. EALL performed the bioinformatic analysis. EALL, AUR wrote the manuscript. EALL, JIF, EZM writing, review, editing, and supervision. JIF, EZM funding acquisition.

## ACKNOWLEDGEMENTS

The authors acknowledge the support from Universidad Nacional del Santa (UNS). They thank Roberto Ferrón, Rolando Quevedo and Milagritos Sánchez from Marinasol for providing the shrimps for the experiment, and Mervin Guevara, Beder Ramírez and Johnny Robles from IMARPE-Tumbes for their support during the development of the experiment.

